# Binless normalization of Hi-C data provides significant interaction and difference detection independently of resolution

**DOI:** 10.1101/214403

**Authors:** Yannick G. Spill, David Castillo, Marc A. Marti-Renom

**Affiliations:** CNAG-CRG, Centre for Genomic Regulation (CRG), Barcelona Institute of Science and Technology (BIST), Baldiri i Reixac 4, 08028 Barcelona, Spain.; Gene Regulation, Stem Cells and Cancer Program, Centre for Genomic Regulation (CRG), Dr. Aiguader 88, 08003 Barcelona, Spain.; Universitat Pompeu Fabra (UPF), Barcelona, Spain.; ICREA, Pg. Lluís Companys 23, 08010 Barcelona, Spain

**Keywords:** Hi-C, normalization, interaction detection, chromatin loops, TADs, Generalized Additive Models, fused lasso

## Abstract

3C-like experiments, such as 4C or Hi-C, have been fundamental in understanding genome organization. Thanks to these technologies, it is now known, for example, that Topologically Associating Domains (TADs) and chromatin loops are implicated in the dynamic interplay of gene activation and repression, and their disruption can have dramatic effects on embryonic development. To make their detection easier, scientists have endeavored into deeper sequencing to mechanically increase the chances to detect weaker signals such as chromatin loops. Part of this mindset can be attributed to the limitations of existing software: the analysis of Hi-C experiments is both statistically and computationally demanding. Here, we devise a new way to represent Hi-C data, which leads to a more detailed classification of paired-end reads and, ultimately, to a new normalization and interaction detection method. Unlike any other, Binless is resolution-agnostic, and adapts to the quality and quantity of available data. We demonstrate its capacities to call interactions and differences and make the software freely available.

## Introduction

Since the invention of 3C-like experiments [1], our perception of the genome has become that of a structured but highly dynamic polymer [2]. In particular, Hi-C experiments [3] made it possible to quantify the frequency of contact between any two locations in the genome. We now know that the mammalian genome of is organized into compartments that, in turn, are partitioned into Topologically Associated Domains (TADs), which holds groups of genes that are preferentially co-expressed. Lately, a series of Hi-C experiments with unprecedented sequencing depth revealed that, at the smallest scale, chromatin loops can form mainly between gene promoters and their enhancers or CTCF bound loci [4]. Yet while it might at first seem that the detection of such events is a mere consequence of better experiments and increased sequencing effort, the computational tool to detect them proves to be crucial. Indeed, the size, noise and complexity of 3C-like experiments also raise completely new research questions for statisticians and computer scientists. A Hi-C experiment on human cells processed at 5 kilobase (kb) resolution yields data comparable to half a million images of 1 megapixel each. Owing to close interdisciplinary collaborations, numerous methods have therefore been developed to post-process 3C-like data [5].

Hi-C experiments [3] produce chimeric DNA fragments, which result from the ligation of two potentially distant parts of the genome that were close in three-dimensional (3D) space. Paired-end sequencing is then used to read the ends of these fragments. Next, a mapper deduces from where, on a reference genome, each end of the read pair comes from. Once paired-end reads have been mapped, Hi-C data is put into matrices at a given resolution. These raw matrices usually show strong systematic biases along both counter-diagonals and rows or columns. It is, therefore, customary to remove these biases through normalization procedures [6–16]. Features, such as TADs [15–21] and chromatin loops [7, 8, 12–14, 16, 22–27] can then be extracted. While cell biologists are in high demand of such algorithms for normalization, TAD and loop calling, there is currently no method that is best for each scenario. Careful and redundant analyses must be made with several tools to conclude the validity of a set of detected interactions.

In this paper, we devise a new way to represent Hi-C data, which casts a different light onto the workings of the Hi-C experiment. By suggesting a new classification of paired-end reads, this representation opens the path to a resolution-agnostic normalization method, coined Binless. This normalization allows to compute matrices that adapt to the size of the features present in the data. We show that in addition to being visually simpler than regular Hi-C maps, they allow for an improved and reproducible interaction detection.

## Results

### Base-resolution view of Hi-C data

A zoomed Hi-C map of the *Caulobacter crescentus* genome [28] at 100 base-pair resolution results in an interesting pattern (Fig. 1A) that prompted us to introduce a new representation of Hi-C data (Fig. 1B). In this new representation, each read is displayed as an arrow in the 2D plane. Projecting the arrow onto the diagonal along the x or y axis, we can retrieve the start, end and orientation of each of the two read pairs (Fig. 1C). Contrary to representing Hi-C data as a matrix of read counts at a given resolution, this base-resolution representation gives insight into the way paired-end reads align around each cut site. Two types of reads can rapidly be distinguished. First, reads that are far from the diagonal are reads with successful re-ligation (or, rarely, mapping errors). Second, those that cluster close to the diagonal correspond to reads in which ligation events were unsuccessful, or which resulted in the re-ligation of the same piece of DNA that was just cut. Depending on their position and orientation relative to a nearby cut site, a new classification can be proposed (Fig. 1D and Supplementary Material). For example, the so-called dangling reads (that is, reads containing fragments of DNA that were digested but not re-ligated) are arrows that stack onto a cut site in this representation.

**Figure 1.**
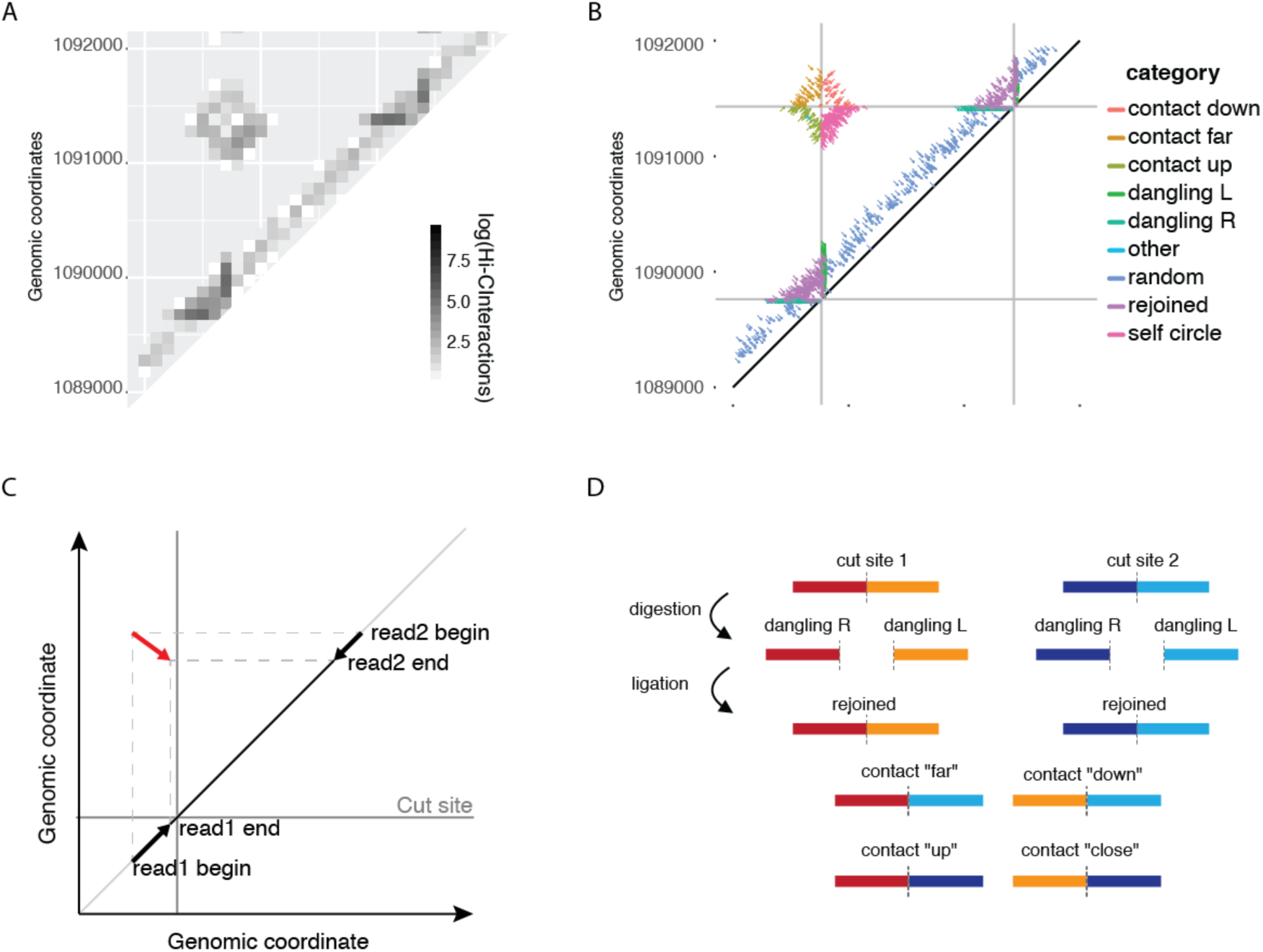
Classification of reads into categories used for Binless normalization. A) Zoomed Hi-C map of *Caulobacter crescentus* at 100bp resolution with the apparent cut-site enrichment of interactions as a square in in the matrix. B) Same data as in panel A now represented as arrows classified for Binless normalization. Vertical and horizontal lines are cut site locations, while diagonal grey line represents the diagonal of the Hi-C matrix. C) Each paired-end read is represented by an arrow. Horizontal and vertical projections on the diagonal reveal the position and direction in which the read was mapped. D) With the new representation, reads are further classified into dangling, rejoined ends and contacts.

This new classification allows to compute three important diagnostics, which are useful for a first evaluation of the quality of the Hi-C experiment. First, the distribution of sonication fragment lengths can be gathered from reads close to the diagonal (Fig. 2A), which can be used to detect problems during the sonication step of the Hi-C protocol. Second, the precise starting points of the dangling ends can also be gathered (Fig. 2B), as they are specific of each restriction enzyme. Spurious peaks in these plots could be indicative of DNA degradation. Third, and most importantly, a local measure of dataset quality can be computed. By assuming that the digestion is 100% efficient, repeated experiments will mainly differ by the efficiency of the ligation step. Therefore, we can examine all reads close to the diagonal (Fig. 1B) and define a ligation ratio (LR) as the number of reads not aligning at a cut site to the total number of reads close to the diagonal. The LR can then be used as a proxy of ligation efficiency. Indeed, even though the LR is a local measure of the Hi-C experiment, it correlates well with the percentage of cis-chromosomal interactions (Fig. 2C), which has been previously used to assess the quality of interaction maps [29]. Importantly, the LR can also be used when the cis-trans ratio cannot be easily calculated or obtained (*i.e.,* partial datasets including those from capture Hi-C).

**Figure 2.**
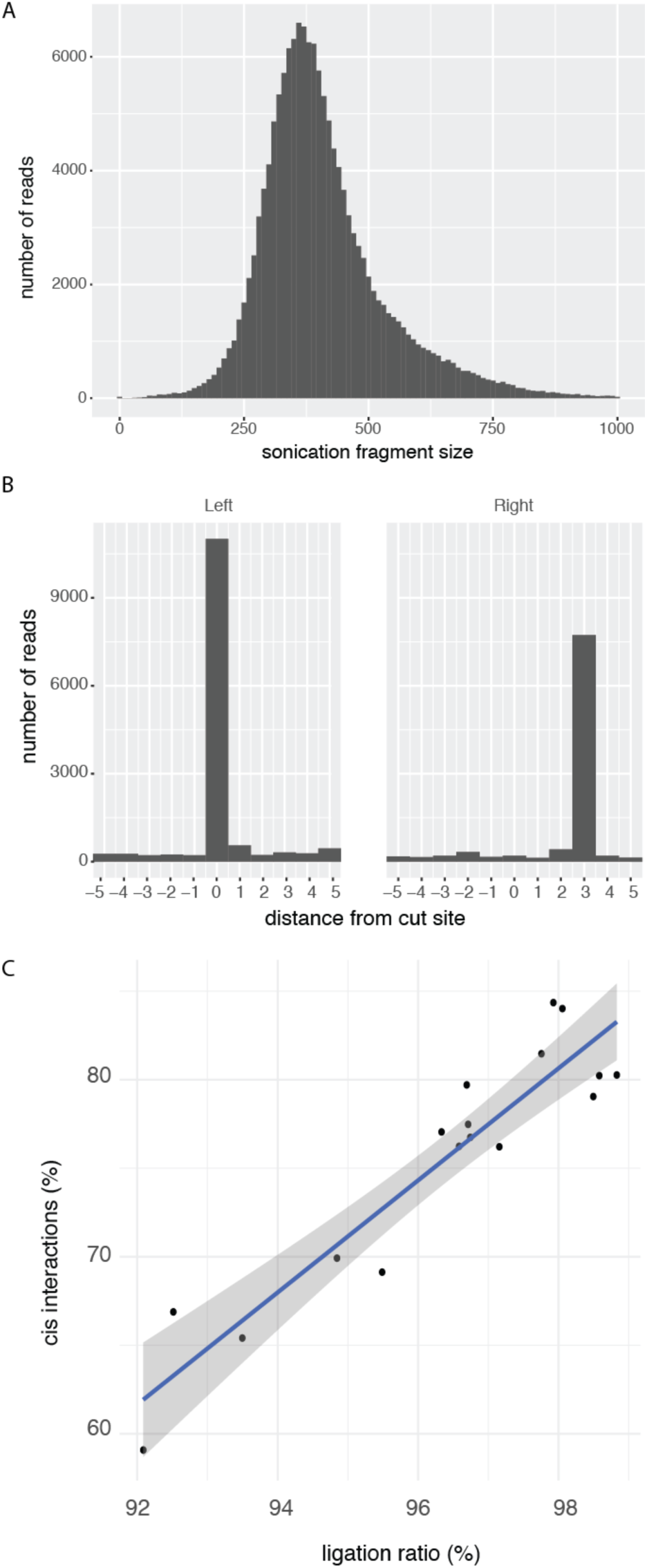
Hi-C diagnostic plots for the PBX1 locus. (Rao [4] GM12878) A) Distribution of the end-to-end distance of dangling ends. In this experiment, the majority of sonication fragments were less than 900 base pairs in size. B) Dangling ends produced by MboI with an overhang of 4 nucleotides. Since by convention, the start of the overhang is at position 0 on the forward strand, it will be at position 3 on the reverse strand. C) Percentage of cis interactions (commonly called “cis/trans ratio”) is correlated with LR score (R^2^ = 0.87, p < 5.8*×*10^−8^). Each point corresponds to one of 18 replicates (HIC001 to HIC018 [4]).

### Binless normalization

The purpose of normalization of 3C-like data is to remove genomic and polymer effects affecting the data. One such bias is observed in the new representation by the proportion of the different dangling ends. It is expected that at constant digestion rate, the number of dangling reads would drop with increased efficiency of ligation at a particular cut site. Here we have estimated these effects using Generalized Additive Model regression [30–32]. This type of regression has non-specific biases, as in iterative correction [7], yet uses a negative binomial count regression framework, similar to HiCNorm [6]. As opposed to iterative correction, however, it does not assume equal visibility of all loci. Crucially, normalization is performed prior to binning the data, as the biases are a property of the data not of the chosen binning. Binless is therefore resolution-independent. Importantly, to assess the biases, it makes use of read pairs that are usually discarded, such as dangling and rejoined reads, which in some cases can represent more than half of the sequencing effort. In addition, it is performed once for all available datasets; that is, if multiple replicates or conditions are available, they are normalized together to maximize the intrinsic information of all datasets.

In Binless, bias removal and signal detection are executed concurrently. For that purpose, a background model capturing local genomic effects and a global distance decay is calculated. On top of this background, a sparse signal, meant to detect significant deviations from the background is calculated (Methods and Supplementary Material). The background regression is done using p-splines [33], whose smoothness adjusts to the quantity of data, and therefore is less prone to over- or under-fitting. The use of smoothing splines is justified when normalizing much sparser Hi-C matrices that that of *Caulobacter crescentus* previously shown, especially if 4 letter cutters are used. For 4-cutters, the number of possible contacts is so large that even very dense datasets, such as the kilobase-resolution datasets of Rao *et al*. [4], only accumulate about 1 contact every 10 cut site intersections (Sup. Fig. 1). To ensure proper normalization and to avoid overfitting, it is therefore essential to share information spatially, which is what Generalized Additive Models were designed for. To show this feature in a genomic application, we took different sub-samplings of the SELP locus. Generalized Additive Model ensured that biases stay as smooth as possible (Sup. Fig. 2).

Binless, which does not make the equal visibility assumption as in the case of ICE normalization, allows row and column sums of a Hi-C matrix to deviate from a reference value. In addition, Binless performs signal detection during normalization. In fact, the simultaneous estimation of normalization biases and signal helps reducing artefacts due to the normalization process (Fig. 3). To demonstrate the advantages of simultaneous normalization and signal detection, we built a synthetic dataset containing a loop and a TAD. We observe that the equal visibility assumption causes long-distance artifacts. Iterative estimation of biases and signal allows the genomic coverage to deviate substantially when signal has been detected, which results in less marked long-range artifacts in the normalized matrices.

**Figure 3.**
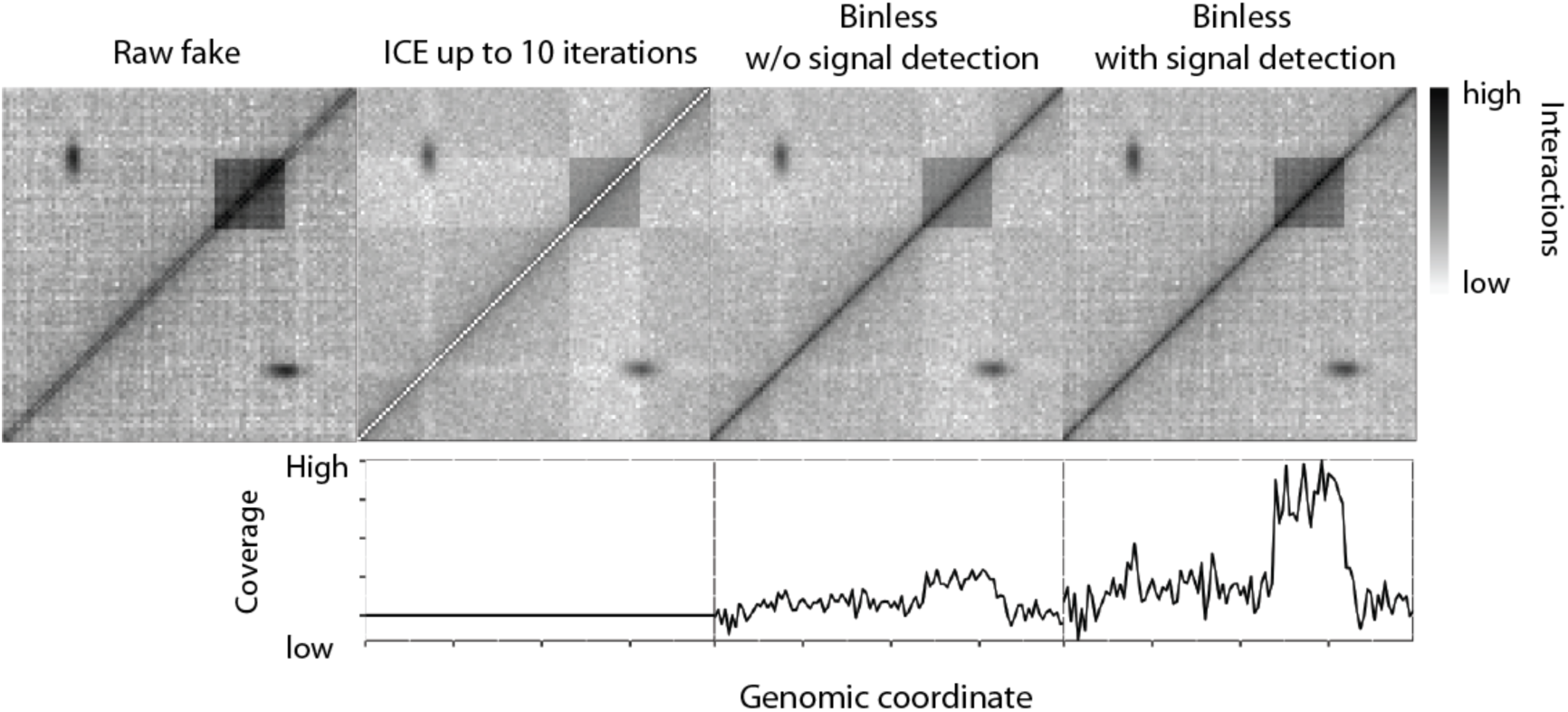
Binless with signal detection performed during normalization removes long-distance artifacts. Fake interaction matrix with genomic decay, noise, a TAD like structure and a loop. From left to right, raw data, ICE normalized data (10 iterations max), Binless normalization with no signal detection, and Binless normalization with signal detection. Bottom graphs show the genomic coverage (column sums) of normalized data for the three approaches.

### Binned detection

Once normalized, the data can be binned at a given resolution (Sup. Fig. 3). As for many normalization methods [7], the normalized matrix is obtained after removing the genomic biases from the raw counts. The resulting matrix has a strong diagonal, as it still contains the decay information. However, to perform interaction detection using a normalized matrix, the polymer effect must be accounted for. Failure to do so would enrich for contacts close to the diagonal. For that purpose, we build the signal matrix, which corresponds to the removal of all estimated biases from the raw counts. The signal matrix informs whether counts in a bin are above or below the estimated background level in that bin or genomic distance. Note that because the normalization procedure is based on a probabilistic model, both normalized and signal matrices come with their corresponding error estimates. These estimates are key for a correct signal detection.

Next, significantly enriched or depleted bins can be called by Bayesian model comparison. We normalized the TBX3 locus on two cell lines (GM12878 and IMR90) by pooling all replicates (Fig. 4A). The calling is performed on each bin of the signal matrix and weighs whether the signal is significantly different from the background at a given threshold, or not (Fig. 4B). Similarly, significant differences with respect to a reference can be called (Fig. 4C), highlighting the appearance of two loops in IMR90 that are not present in GM12878. Note that datasets can be grouped, e.g., by condition, to improve the power of the detection.

**Figure 4.**
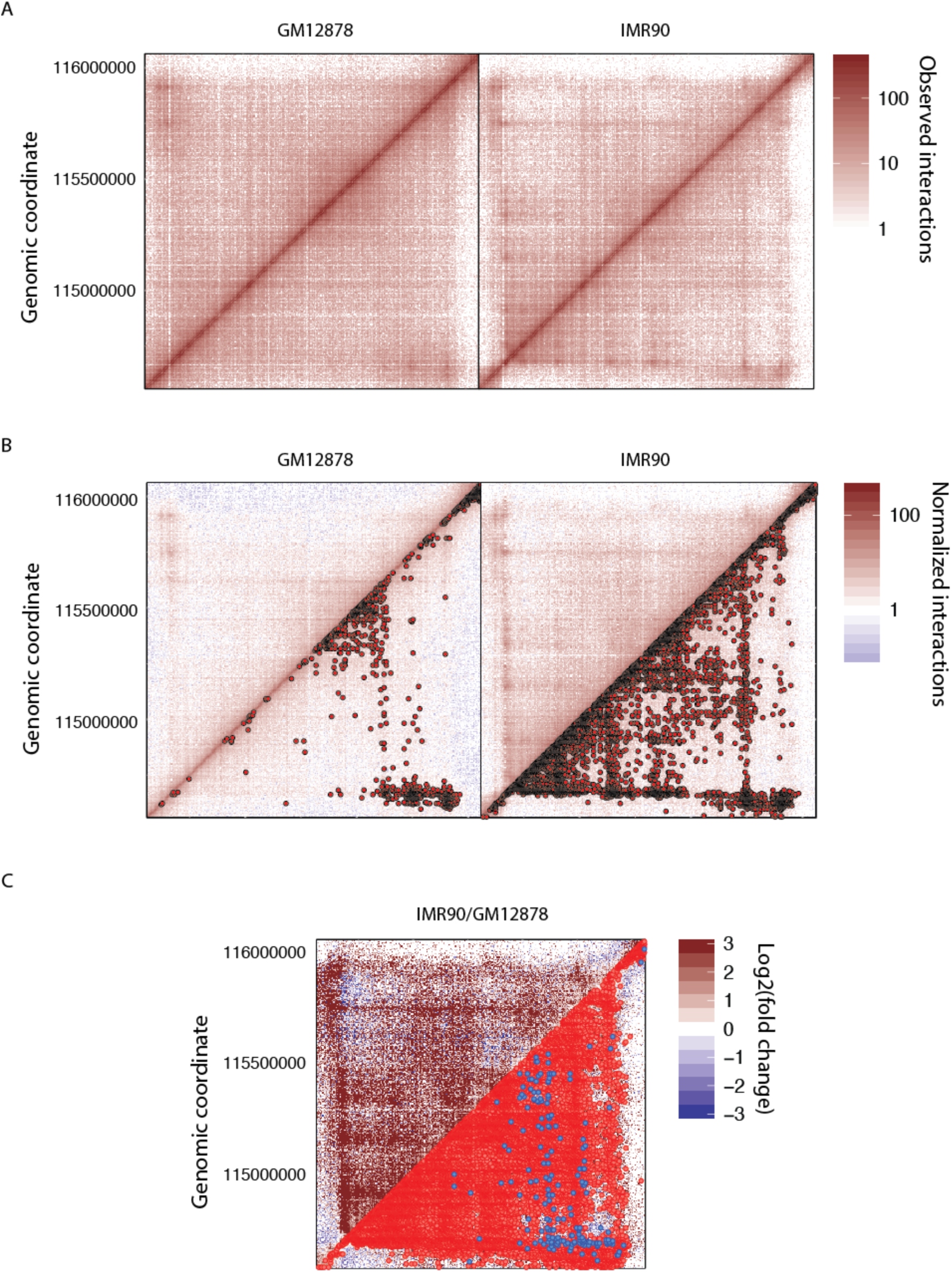
Bin-wise interaction detection on the TBX3 locus in the human chromosome 12. A) Raw counts (left: GM12878, right: IMR90, data from [4]). B) Normalized matrices overlaid with statistically significant interactions. Only the first 1000 significant interactions are shown. C) Bin-wise differential signal between IMR90 and GM12878 cell lines. Red and blue dots are significantly enriched and depleted in IMR90 compared to GM12878, respectively.

### Binned detection lacks sensitivity

A number of problems arise in all binned interaction detection methods, and the one proposed here is no exception. The significance of interactions depends on the chosen resolution. This was already observed previously and is a consequence of the binning since loops are usually called at 1-10kb, TADs at 50-100kb and compartments at 100-1000kb resolution [34]. The best resolution at which to call interactions is an open question, and depends on data quality. Importantly, at a given resolution, the number of common interactions between replicates is normally low, as was shown recently in a comparison of all existing interaction detection tools for Hi-C [34]. We normalized the TBX3 locus on two cell lines (GM12878 and IMR90) with each two pools of replicates (Sup. Fig. 4). Interaction detection results in an overlap which is low (Fig. 4C). Most importantly, there can be less interactions in common between replicates than between cell lines, irrespective of the resolution of the calling. Difference detection between two pools of replicates also reports false interactions (Fig. 4D). However, it can be noted qualitatively that most features, such as loops and TADs, have been captured by the interaction detection. Indeed, we can see that significant interactions are called more often around these regions than elsewhere. A better accounting of the spatial dependence of interactions is, as we will now show, the key to an improved interaction and difference detection.

### Binless detection

A binless matrix is a matrix whose bins adapt to the size of the features to be detected. This adaptation is made using Fused Lasso regression [35–37]. It is possible to do this adaptation while maintaining statistical significance. This is because the normalization we presented and used prior to peak detection is resolution-agnostic as well. Fused lasso regression builds on a high-resolution matrix representation of the data (called its base resolution), and fuses neighboring bins if they are estimated to have a common signal. It is an alternative approach to other neighborhood filtering approaches [38, 39], for which highly efficient implementations are available [40–45]. Binless interaction and difference detection using Fused Lasso regression addresses the problems discussed above. It has two key advantages over interaction detection at a fixed resolution. First, spurious peaks are removed, which makes the matrices visually easier to interpret (Fig. 5A,B). Second, this adaptive fusion causes a dramatic increase in detection sensitivity since it allows simultaneous detection of both sharp and strong features, such as chromatin loops, along with larger but weaker features, such as TADs. Crucially, significant differences are much more sensitive. For example, four loops are significantly enriched in IMR90 cells compared to GM12878 (Fig. 5C). At the same time, if we compare replicate pools to each other (Fig. 6), no significant differences were observed between two pools of GM12878 replicates (Fig. 6C). In contrast, binned interaction detection methods normally report significant differences between replicates, and sometimes even larger differences than between two different experimental conditions (see e.g. [34] and Sup. Fig. 4D). Because Binless interaction detection works by sharing spatial information optimally, such a high false positive rate is less likely to appear. In fact, on 8 different loci in two different cell types, there are no false discoveries using the Binless difference detection, while up to 5% of false discoveries were made with binned difference detection (Table 1). Binless matrices report a result that is both strikingly more visual and sensitive, as we show in three loci (Fig. 7). Among others, in ADAMTS1 (Fig. 7A,B), six loops are significantly enriched in IMR90 and three TADs appear. In FOXP1 (Fig 7C,D), we detect an enrichment of interactions within the central TAD. In SEMA3C (Fig. 7E,F), a TAD is split in two, and an additional loop appears.

**Table 1.** Percentage of significant differences between two replicates of the same experiment, for 16 comparisons on 8 loci and 2 cell lines.

| resolution | Binned |  |  | Binless |  |  |
| --- | --- | --- | --- | --- | --- | --- |
|  | min (%) | average (%) | max (%) | min (%) | average (%) | max (%) |
| 5000 | 0 | 0.12 | 1.05 | 0 | 0 | 0 |
| 10000 | 0 | 0.74 | 5.40 | 0 | 0 | 0 |

**Figure 5.**
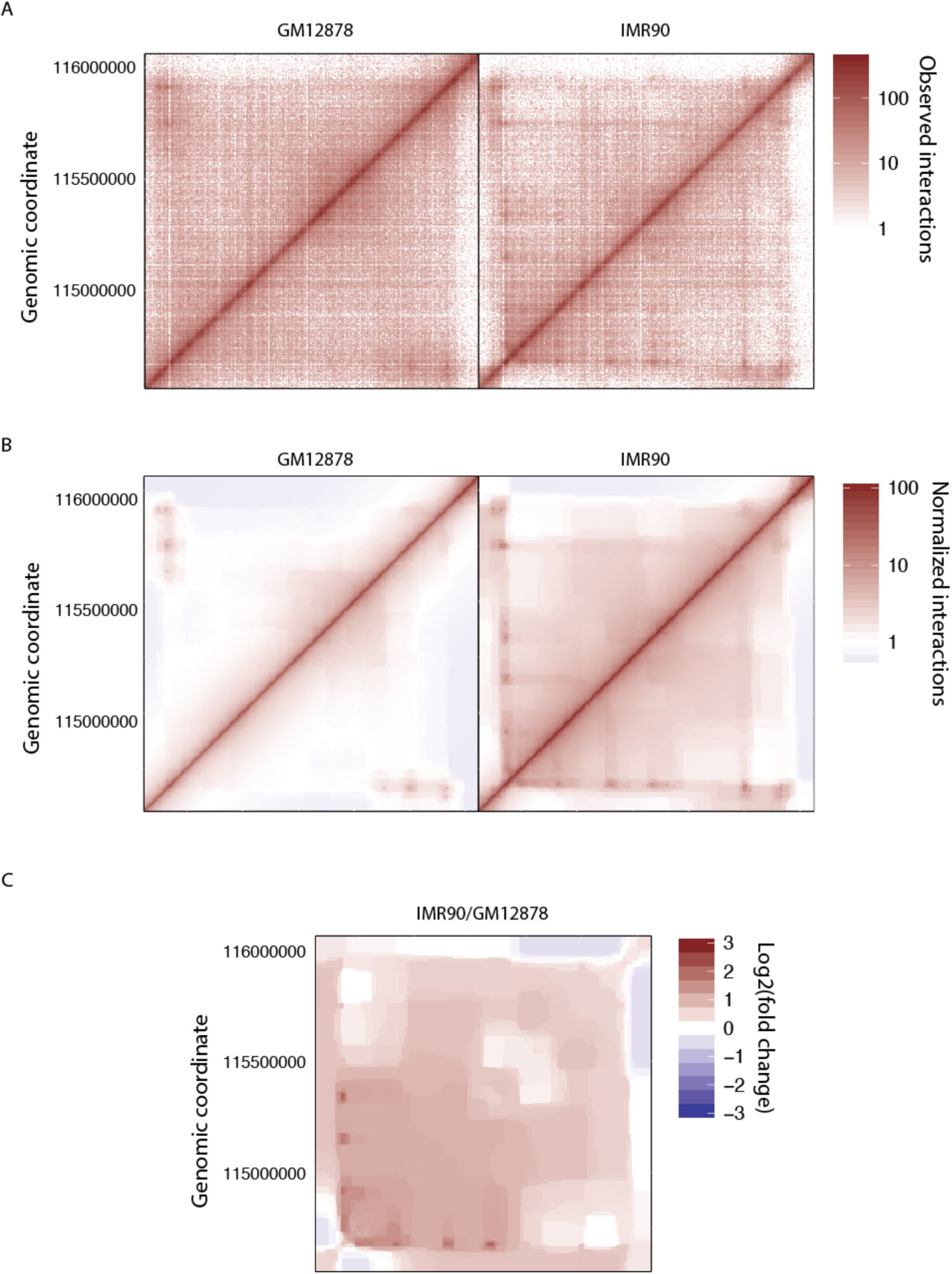
Binless interaction and difference detection on the TBX3 locus. From left to right: GM12878 all datasets, IMR90 all datasets, and differences between IMR90 and GM12878.

**Figure 6.**
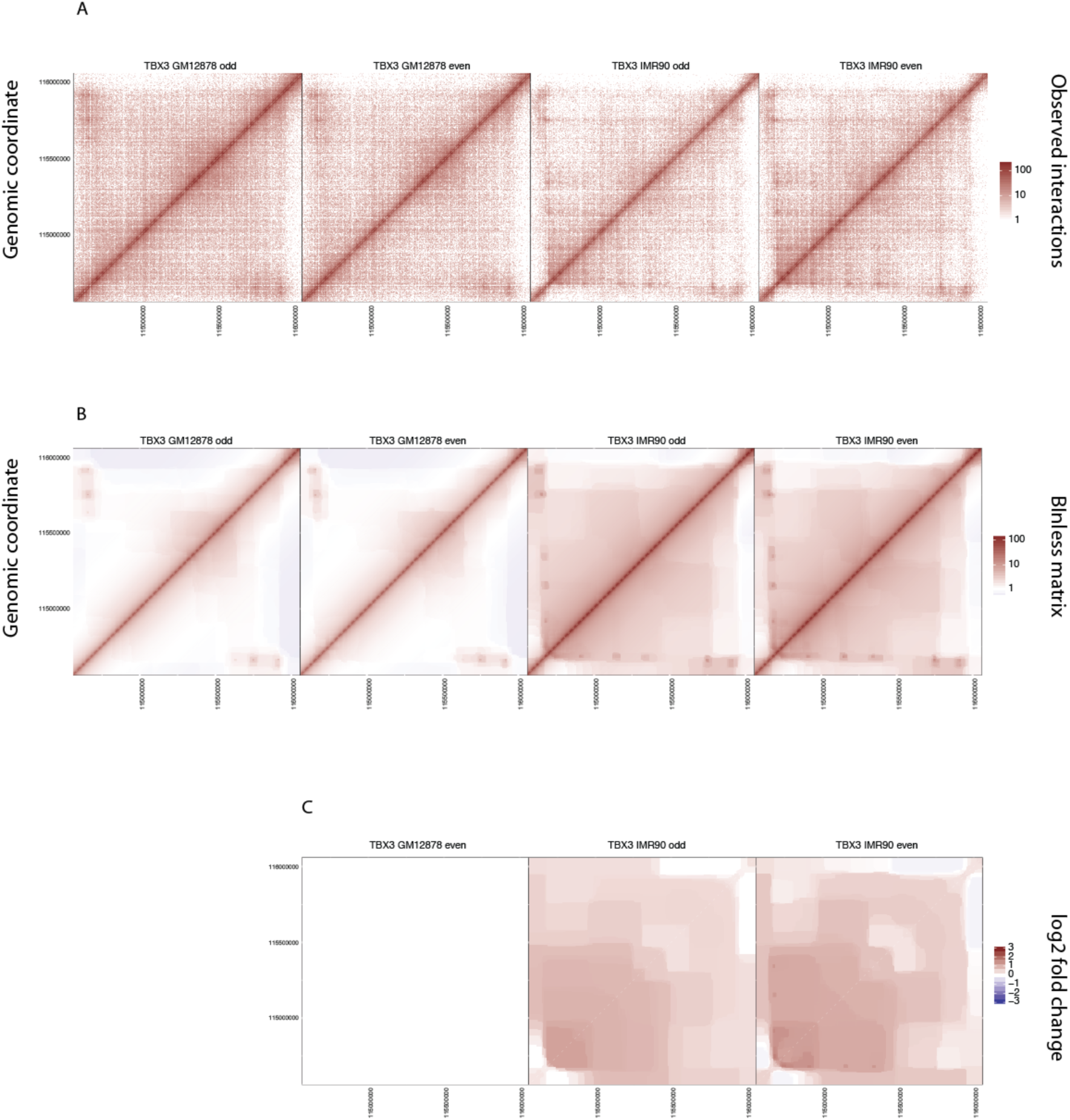
Interaction detection on the TBX3 locus in a paired setting. (Rao [4] GM12878 replicates HIC001-18 and IMR90 replicates HIC050-56 pooled in two groups of even- or odd-numbered replicates). A) Differential interaction maps at 10Kb bins and statistically significant differences in the indicated replicate with respect to GM12878 even replicate numbers. On the left of the panel are the number of interactions in each dataset, the number of common interactions between datasets and the Jaccard index (percent of common interactions). B) Same as panel A but maps were generated with bins at 5Kb resolution. C) Same as panel B for binless signal and sample as well as differential matrices.

**Figure 7.**
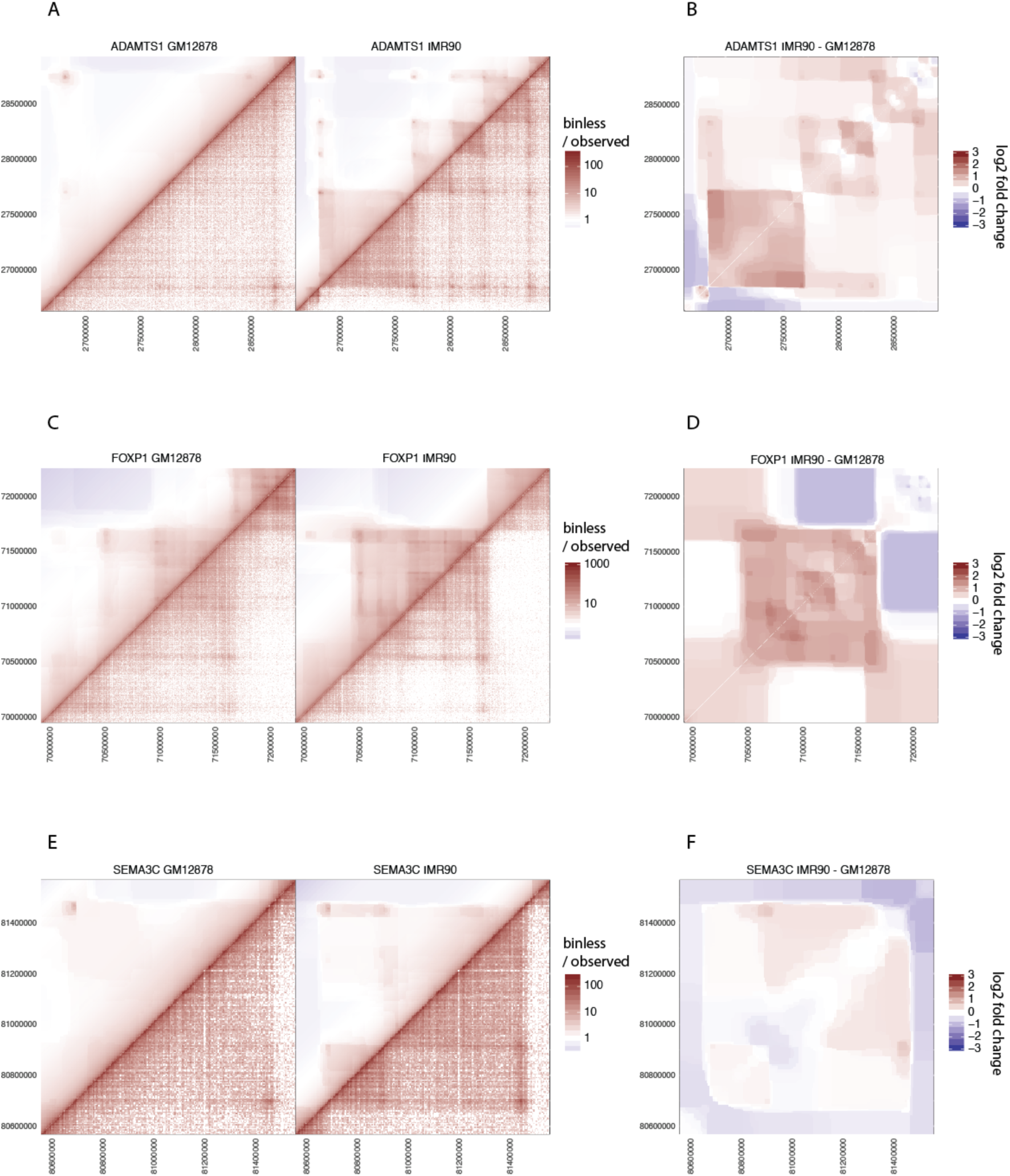
Binless interaction and difference detection captures significant changes in loops and TADs. Raw data and binless matrices (A, C, E) and significant differences (B, D, F) in three different datasets (A,B: ADAMTS1; C,D: FOXP1; SEMA3C: E,F).

## Discussion

In this paper, we present Binless, a method to normalize 3C-like data in a robust, resolution-independent and statistically significant way. Two types of strategies exist to normalize Hi-C data, as was reviewed recently [5]. On one hand, explicit methods assume that all biases affecting Hi-C data are known and can be provided as input to the normalization software; for example, HiCNorm [6] requires three genomic tracks for GC content, mappability and fragment length. On the other hand, implicit methods make one strong theoretical assumption on Hi-C matrices, such as equal visibility of all loci in ICE [7]. They then deduce the biases that must be subtracted to recover normalized Hi-C matrices. Our binless normalization uses the negative binomial regression framework that proved its validity in HiCNorm [6]. However, we feel that the knowledge of the true sources of bias in 3C-like data is still limited. Relying only on GC content, mappability and fragment length is too approximate, although it clearly captures the main effects. We therefore follow the path of implicit methods and estimate the biases only based on the Hi-C data. We however do not make the equal visibility assumption. Instead, we propose that all loci be equally visible on average only, and perform signal detection and normalization simultaneously. If a locus conflicts with its immediate neighbors, then its visibility should be allowed to deviate from the average (Fig. 3).

As was noted recently [5], the resolution of a Hi-C matrix is often chosen based on the distance between two loci of interest. Indeed, it is expected that higher resolution can be reached for closer loci. Sequencing depth then dictates where to fix the tradeoff between high resolution and maximum distance. With Binless, it is possible to perform normalization, interaction and difference detection entirely without specifying a Hi-C matrix resolution. Internally, Binless does not simply pick an optimal resolution based on some heuristic. The resolution is adaptive, and depends on the position in the Hi-C matrix. Therefore, there is no tradeoff to make. The fused lasso algorithm used for that purpose ensures that, at each position, the local bin size is neither too big, which could lead to averaging out some features, nor too small, which would increase the noise. Incidentally, Binless is able to detect loops and TADs within the same matrix.

In the past, resolution, as in other fields such as cryo-electron microscopy, has been associated with quality of the data. In this paper, however, we prove that it is possible and advantageous to normalize Hi-C data in a resolution-agnostic way, using Binless matrices. Therefore, how can the quality of a dataset be assessed? We propose two solutions. First, detection power scales with the number of ligated reads. The ligation ratio (LR, Fig. 2) can be used to rate the quality of a library. The LR is a local measure of ligation efficiency, and can be evaluated at low cost, since only a few thousand reads are typically necessary to compute it. Second, binless matrices have a base resolution, which can be seen like the pixel size of a detector. Binless patches vary substantially in size (Fig. 8), and as expected, the larger the patch, the more reads it accumulates. However, because the fused lasso algorithm adapts to the features it detects, the resulting bins have an approximately constant read density, independent of patch size. We therefore propose to use this average read density per patch as a proxy for dataset quality.

**Figure 8.**
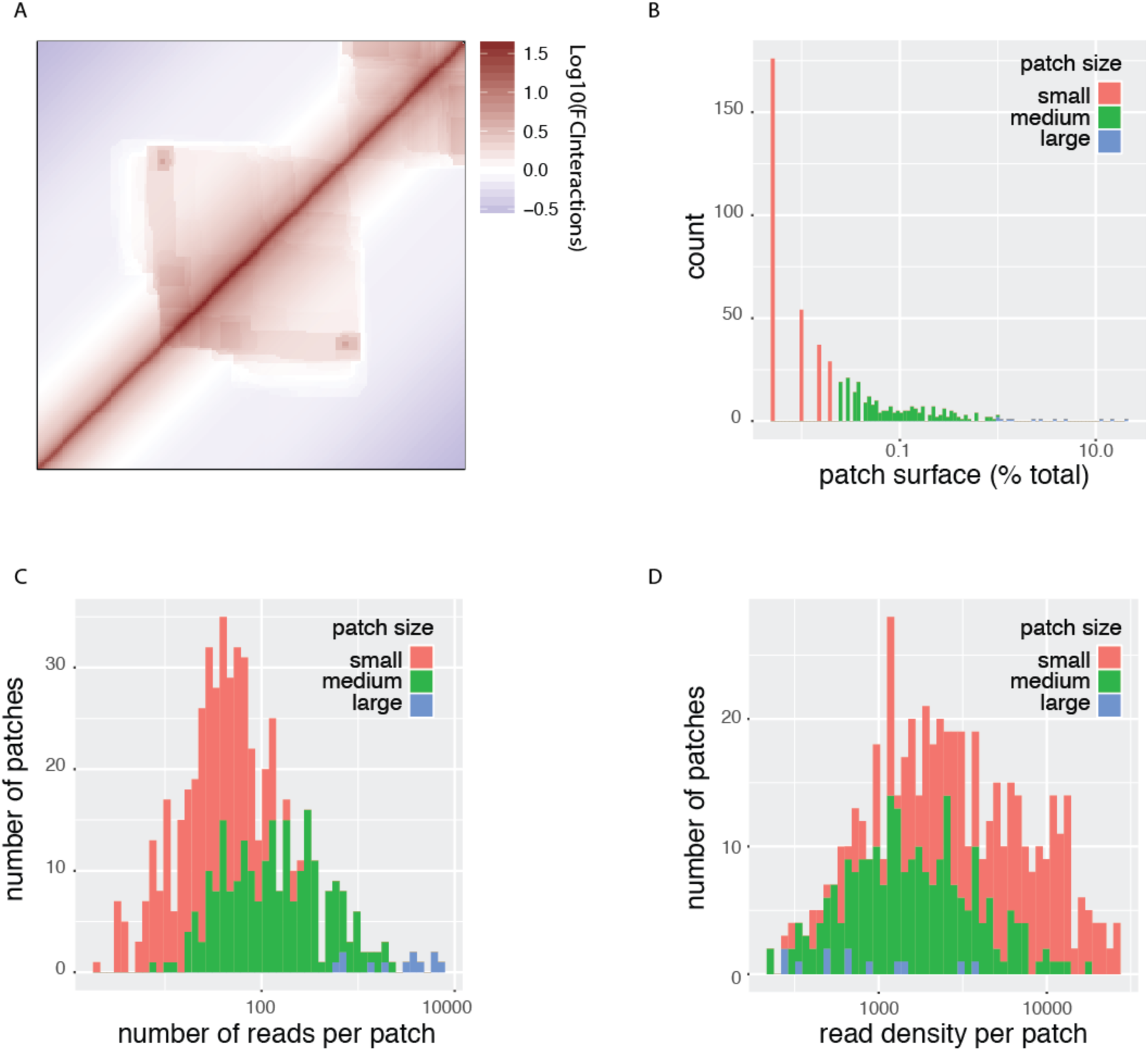
Binless patch statistics for an 1Mb example region (HIC003, hg19 [4], called “Replicate H” in [34]). A) binless matrix of the region. B) Distribution of patch sizes in the binless matrix. C) Distribution of the number of reads per patch. D) Distribution of the read density per patch.

## Methods

### Base-resolution view of Hi-C data

Paired-end reads are processed using the TADbit pipeline [46]. The input to Binless is the reads intersection file. It contains the genomic location, length and strand for both ends of each read, as well as the coordinates of the closest upstream and downstream cut sites. It is assumed that the first read is always upstream of the second read. Duplicate reads are removed when reading the inputs. At this step, sonication fragment length and dangling end positions should be provided by the user.

These reads are then classified as shown schematically in Fig. 1. We define several categories. A left or right “dangling read” is a DNA molecule that starts or ends, respectively, on a cut-site, with both ends mapping on opposite strands. A “rejoined read” spans across a restriction site. The name is because it has likely been re-ligated. A “self-circle” corresponds to ligation of the two ends of a fragment. “Random reads” align close to the diagonal and on the same fragment, and point towards the diagonal in the base-resolution representation. They are thought to be genomic DNA, and are not specific to the Hi-C experiment. Most importantly, there are four “contact types”, depending on which quadrant they can be found in. Up and down contacts are such that both read ends align upstream and downstream, respectively, of the closest restriction fragment. Close and far contacts are closer or further, respectively, from the diagonal than the intersection of their cut sites. All contact types must point towards the restriction intersection in the base-resolution representation. Note that for neighboring cut sites, self-circles replace the close contact category. Finally, reads that cannot be classified (because they are too far from a restriction site, or because their direction does not match) are put in “other” category. Proportions in each category are shown for four datasets (Sup. Table 1).

The quantities in Fig. 2 were computed as follows. Cis-trans ratio: Number of filtered reads arising within chr1 divided by total number of filtered reads with one end mapping on chr1. Filtering options are those used by default in TADbit. LR: Select all reads whose end-to-end distance mapped at most at 900bp from each other, on the PBX1 locus (Rao [4] GM12878). Further, keep only reads where read 1 aligns on the forward strand, while read 2 aligns on the reverse (assuming read 1 is upstream of read 2). Note the position of dangling ends (Fig. 2). The LR is the number of reads that do not start or end at a cut site, divided by the total number of reads.

### Normalization

<u>Exact model</u>. The negative binomial regression we employ has likelihoods of the following form (see Supplementary Material for a complete overview)

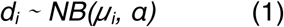

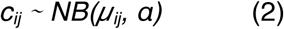

where *d*_*i*_ is the number of dangling or rejoined reads at cut site *i* and *c*_*ij*_ is the number of reads in one of the four contact categories, observed between cut sites *i* and *j*. *µ*_*i*_ and *µ*_*ij*_ are the respective means, to be estimated, and *α* the dispersion parameter of the negative binomial. The means *µ*_*ij*_ are parametrized using three “background” splines *ι*, *ρ* and *f*, and one “signal” term s. The *µ*_*i*_ are parametrized only using only the first two splines, as we now explain. The efficiency of detection of a particular contact has been shown to be decomposable into genome-specific biases for each of the two reads in a read pair [7]. For reads aligning to the left (respectively right) of a cut site *i*, the number of contacts involving *i* are therefore made proportional to their genomic bias *ι*_*i*_ (respectively *ρ_i_*). *ι* and *ρ* are modelled using p-splines [30, 32, 33, 47–49]. The polymer nature of chromatin is thought to make Hi-C contact probabilities decrease with the genomic distance between two cut sites. Therefore, the number of contacts involving cut sites *i* and *j* are made proportional to the decay bias *f*_*ij*_, which is forced to decrease with the genomic distance between *i* and *j*. We use a smooth constrained additive model for *f* [31]. When the ligation efficiency for a cut-site decreases, one can expect a depletion in the number of contacts and an enrichment of dangling ends. Therefore, dangling ends are made to follow the opposite trend of the counts, and are biased by *ι* for left-dangling and *ρ* by right-dangling ends. Rejoined ends follow the (geometric) average bias at this cut site. Finally, a sparse 2D term *s*_*ij*_ is meant to fit the signal that departs significantly from the background modelled by the genomic and polymer effects. This term is modelled using the sparse 2D generalized fused lasso on a triangle grid graph [36, 41, 43–45].

<u>Fast normal approximation.</u> Ideally, all parameters are optimized together. However only small datasets (less than about 100 cut sites) can be normalized in this way. For larger datasets, we resort to a fast and approximate coordinate descent algorithm. In a nutshell, instead of optimizing all parameters at once, we optimize parameters relevant to genomic biases, diagonal decay, dispersion (using Stan [50, 51]) and signal (using gfl [45]) separately and iteratively. In each separate optimization, we compute the biases not using the individual counts, but using weighted average log-counts. This grouping by rows, counter diagonals or signal bins is what allows the computation to be orders of magnitude faster. Grouping is made possible by a repeated normal approximation to the log likelihood of the counts. This approximation, known as Iteratively Re-weighted Least Squares (IRLS) is very common in all types of generalized regressions [32, 52]. The estimation of the dispersion is done differently. We take all rejoined and dangling read counts, and sample a large (default 10^5^) number of counts, including zeros, from the data. We then estimate the dispersion using a simple Stan model [50, 51].

<u>Available outputs</u>. Once several datasets have been normalized together, a number of matrices can be produced at any resolution. The signal matrix corresponds to removing all estimated background from the observed data. The normalized matrix corresponds to removing all genomic biases. Each matrix comes with corresponding error estimates, that are provided using the IRLS approximation (Supplementary Material). For the signal matrices, significant interactions are detected using Bayesian Model comparison. Interactions are deemed significant if *K/(1 + K)* is larger than a threshold (default 0.95), where *K* is the Bayes factor of a model with signal with respect to a model without. Similarly, significant differences to a reference are computed by comparing a model with two different signals to a model with only one common signal.

Binless signal matrices are the signal term obtained during normalization. They can also be recomputed at a different base resolution afterwards. Their unit is a fold change with respect to the background. Because sparsity was enforced while estimating the signal, the resulting matrix is nonzero when the signal is statistically significant. The binless signal matrix can be shown with an added decay bias. Such a matrix, which we simply call binless matrix, is visually closer to the raw data, but its unit is a fold change with respect to a background without polymer decay. Finally, binless differences between datasets can be computed, and their unit is a fold change between one dataset and a reference. All aforementioned matrices can be grouped (e.g. by condition) to improve the detection sensitivity.

<u>Recommendations.</u> In designing Binless, we attempted to minimize the number of free parameters. Yet, some of them are left to the choice of the user. As a general rule, their choice should not impact the resulting normalization. For example, the number of iterations should be large enough to reach convergence of the algorithm, which can be monitored using diagnostic plots Binless provides. Most importantly, the number of basis functions per kilobase controls the maximum wiggliness of the genomic biases. If it is large, the computational burden is high and the normalization can, for very large datasets, become unstable. If it is too small, the genomic biases will not be estimated properly. We suggest to start with a value of 30 and increase it if necessary. Similarly, for binless detection, the base resolution should be as small as the smallest feature one hopes to detect. Out of computational considerations, we recommend a base resolution of 5kb. Once normalized, the data can be rebinned if necessary.

### Figure generation

All data pertaining to datasets used in this article can be found in Sup. Table 2. All data was processed with the TADbit pipeline [46] and binless 0.9.0. Binless takes as input the mapped pair-end reads of each dataset. We used 30 basis functions per kb and a base resolution of 10kb. For subsampling of the data in Sup. Fig. 2, we took a subset of all available reads by drawing the read count from a binomial distribution (coin tossing). Each dataset was then normalized independently, without signal modelling.

### Code availability

Binless is an R/C++ package using Stan [50, 51] and gfl [45] and is available at github.com/3DGenomes/binless

## Acknowledgements

The authors are grateful to François Le Dily, Guillaume J. Filion, Francesca Di Giovanni, Simon Heath, Emanuele Raineri, François Serra and Enrique Vidal for fruitful discussion.

## Author contributions

Y.G.S. and M.A.M-R. designed the method. Y.G.S. developed the method and analyzed the Hi-C datasets. Y.G.S. and D.C. implemented the package. Y.G.S and M.A.M-R. wrote the manuscript.

## References

[1] Job Dekker, Karsten Rippe, Martijn Dekker, and Nancy Kleckner. “Capturing Chromosome Conformation”. In: Science 295.5558 (2002), pp. 1306–1311. doi: 10.1126/science.1067799.

[2] Job Dekker, Marc A. Marti-Renom, and Leonid A. Mirny. “Exploring the three-dimensional organization of genomes: interpreting chromtin interaction data”. In: Nature Rev. Genet. 14 (2013), pp. 390–403.

[3] Erez Lieberman-Aiden, Nynke L. van Berkum, Louise Williams, Maxim Imakaev, Tobias Ragoczy, Agnes Telling, Ido Amit, Bryan R. Lajoie, Peter J. Sabo, Michael O. Dorschner, Richard Sandstrom, Bradley Bernstein, M. A. Bender, Mark Grou-dine, Andreas Gnirke, John Stamatoyannopoulos, Leonid A. Mirny, Eric S. Lander, and Job Dekker. “Comprehensive Mapping of Long-Range Interactions Reveals Folding Principles of the Human Genome”. In: Science 326.5950 (2009), pp. 289–293. doi: 10.1126/science.1181369.

[4] Rao S. S. P., Huntley M. H., Durand N. C., Stamenova E. K., Bochkov I. D., Robinson J. T., Sanborn A. L., Machol I., Omer A. D., Lander E. S., and Lieberman Aiden E. “A 3D map of the human genome at kilobase resolution reveals principles of chromatin looping”. In: Cell 159 (7 2014), pp. 1665–1680. doi: dx.doi.org/10.1016/j.cell.2014.11.021.

[5] Anthony D. Schmitt, Ming Hu, and Bing Ren. “Genome-wide mapping and analysis of chromosome architecture”. In: Nat Rev Mol Cell Biol 17 (12 2016), pp. 743–755.

[6] Hu M, Deng K, Selvaraj S, Qin Z, Ren B, and Liu JS. “HiCNorm: removing biases in Hi-C data via Poisson regression”. In: Bioinformatics 28.23 (2012), pp. 3131–3133. doi: 10.1093/bioinformatics/bts570.

[7] Imakaev M, Fudenberg G, McCord RP, Naumova N, Goloborodko A, Lajoie BR, Dekker J, and Mirny LA. “Iterative correction of Hi-C data reveals hallmarks of chromosome organization”. In: Nature Methods 9 (2012), pp. 999–1003. doi: 10.1038/nmeth.2148.

[8] S. Heinz, C. Benner, N. Spann, E. Bertolino, Y. C. Lin, and P. Laslo. “Simple combinations of lineage-determining transcription factors prime cis-regulatory elements required for macrophage and B cell identities”. In: Mol Cell 38 (2010). doi: 10.1016/j.molcel.2010.05.004.

[9] Nicolas Servant, Bryan R. Lajoie, Elph ge P. Nora, Luca Giorgetti, Chong-Jian Chen, Edith Heard, Job Dekker, and Emmanuel Barillot. “HiTC: exploration of high-throughput C experiments”. In: Bioinformatics 28.21 (2012), pp. 2843–2844. doi: 10.1093/bioinformatics/bts521.

[10] W. Li, K. Gong, Q. Li, F. Alber, and X. J. Zhou. “Hi-Corrector: a fast, scalable and memory-efficient package for normalizing large-scale Hi-C data”. In: Bioinformatics 31 (2015). doi: 10.1093/bioinformatics/btu747.

[11] M. E. Sauria, J. E. Phillips-Cremins, V. G. Corces, and J. Taylor. “HiFive: a tool suite for easy and efficient HiC and 5C data analysis”. In: Genome Biol 16 (2015). doi: 10.1186/s13059-015-0806-y.

[12] G. Castellano, F. Dily, A. Hermoso Pulido, M. Beato, and G. Roma. “Hi-Cpipe: a pipeline for high-throughput chromosome capture”. In: bioRxiv (2015).

[13] N. Servant, N. Varoquaux, B. R. Lajoie, E. Viara, C. -. J. Chen, and J. -. P. Vert. “HiC-Pro: an optimized and flexible pipeline for Hi-C data processing”. In: Genome Biol 16 (2015). doi: 10.1186/s13059-015-0831-x.

[14] M. W. Schmid, S. Grob, and U. Grossniklaus. “HiCdat: a fast and easy-to-use Hi-C data analysis tool”. In: BMC Bioinf 16 (2015). doi: 10.1186/s12859-15-0678-x.

[15] F. Serra, D. Baù, G. Filion, and M. A. Marti-Renom. “Structural features of the fly chromatin colors revealed by automatic three-dimensional modeling”. In: bioRxiv (2016).

[16] Charalampos Lazaris, Stephen Kelly, Panagiotis Ntziachristos, Iannis Aifantis, and Aristotelis Tsirigos. “HiC-bench: comprehensive and reproducible Hi-C data analysis designed for parameter exploration and benchmarking”. In: BMC Genomics 18.1 (2017), p. 22. doi: 10.1186/s12864-16-3387-6.

[17] Jesse R. Dixon, Siddarth Selvaraj, Feng Yue, Audrey Kim, Yan Li, Yin Shen, Ming Hu, Jun S. Liu, and Bing Ren. “Topological domains in mammalian genomes identified by analysis of chromatin interactions”. In: Nature 485 (7398 2012), pp. 376–380. doi: 10.1038/nature11082.

[18] Celine L vy-Leduc, M. Delattre, T. Mary-Huard, and S. Robin. “Two-dimensional segmentation for analyzing Hi-C data”. In: Bioinformatics 30.17 (2014), pp. i386–i392. doi: 10.1093/bioinformatics/btu443.

[19] Darya Filippova, Rob Patro, Geet Duggal, and Carl Kingsford. “Identification of alternative topological domains in chromatin”. In: Algorithms for Molecular Biology 9.1 (2014), p. 14. doi: 10.1186/1748-7188-9-14.

[20] Emily Crane, Qian Bian, Rachel Patton McCord, Bryan R. Lajoie, Bayly S. Wheeler, Edward J. Ralston, Satoru Uzawa, Job Dekker, and Barbara J. Meyer. “Condensin-driven remodelling of X chromosome topology during dosage compensation”. In: Nature 523 (7559 2015), pp. 240–244. doi: 10.1038/nature14450.

[21] Caleb Weinreb and Benjamin J. Raphael. “Identification of hierarchical chromatin domains”. In: Bioinformatics 32.11 (2016), pp. 1601–1609. doi: 10.1093/ bioinformatics/btv485.

[22] Ferhat Ay, Timothy L. Bailey, and William Stafford Noble. “Statistical confidence estimation for Hi-C data reveals regulatory chromatin contacts”. In: Genome Research 24.6 (2014), pp. 999–1011. doi: 10.1101/gr.160374.113.

[23] S Wingett, P Ewels, M Furlan-Magaril, T Nagano, S Schoenfelder, P Fraser, and S Andrews. “HiCUP: pipeline for mapping and processing Hi-C data [version 1; referees: 2 approved, 1 approved with reservations]”. In: F1000Research 4.1310 (2015). doi: 10.12688/f1000research.7334.1.

[24] Y. -. C. Hwang, C. -. F. Lin, O. Valladares, J. Malamon, P. P. Kuksa, and Q. Zheng. “HIPPIE: a high-throughput identification pipeline for promoter interacting enhancer elements”. In: Bioinformatics 31 (2015). doi: 10.1093/bioinformatics/ btu801.

[25] Aaron T.L. Lun and Gordon K. Smyth. “diffHic: a Bioconductor package to detect differential genomic interactions in Hi-C data”. In: BMC Bioinformatics 16.1 (2015), p. 258. doi: 10.1186/s12859-015-0683-0.

[26] Borbala Mifsud, Filipe Tavares-Cadete, Alice N Young, Robert Sugar, Stefan Schoenfelder, Lauren Ferreira, Steven W Wingett, Simon Andrews, William Grey, Philip A Ewels, Bram Herman, Scott Happe, Andy Higgs, Emily LeProust, George A Follows, Peter Fraser, Nicholas M Luscombe, and Cameron S Osborne. “Mapping long-range promoter contacts in human cells with high-resolution capture Hi-C”. In: Nat Genet 47 (6 2015), pp. 598–606. doi: 10.1038/ng.3286.

[27] Stefan Schoenfelder, Mayra Furlan-Magaril, Borbala Mifsud, Filipe Tavares-Cadete, Robert Sugar, Biola-Maria Javierre, Takashi Nagano, Yulia Katsman, Moorthy Sakthidevi, Steven W. Wingett, Emilia Dimitrova, Andrew Dimond, Lucas B. Edelman, Sarah Elderkin, Kristina Tabbada, Elodie Darbo, Simon An-drews, Bram Herman, Andy Higgs, Emily LeProust, Cameron S. Osborne, Jennifer A. Mitchell, Nicholas M. Luscombe, and Peter Fraser. “The pluripotent regulatory circuitry connecting promoters to their long-range interacting elements”. In: Genome Research 25.4 (2015), pp. 582–597. doi: 10.1101/gr.185272.114.

[28] Tung B. K. Le, Maxim V. Imakaev, Leonid A. Mirny, and Michael T. Laub. “High-Resolution Mapping of the Spatial Organization of a Bacterial Chromosome”. In: Science 342.6159 (2013), pp. 731–734. doi: 10.1126/science. 1242059.

[29] Lajoie BR, Dekker J, and Kaplan N. “The Hitchhiker’s guide to Hi-C analysis: Practical guidelines”. In: Methods 72 (2015), pp. 65–75. doi: 10.1016/j.ymeth. 2014.10.031.

[30] Hastie T. and Tibshirani R. “Generalized Additive Models”. In: Stat. Sci. 1 (3 1986), pp. 297–318.

[31] Pya N. and Wood S. N. “Shape constrained additive models”. In: Stat. Comput. 25 (3 2015), pp. 543–559.

[32] S.N Wood. Generalized Additive Models: An Introduction with R. Chapman and Hall/CRC, 2006.

[33] Eilers P. H. C. and Marx B. D. “Flexible Smoothing with B-splines and penalties”. In: Stat. Sci. 11 (2 1996), pp. 89–102.

[34] Mattia Forcato, Chiara Nicoletti, Koustav Pal, Carmen Maria Livi, Francesco Ferrari, and Silvio Bicciato. “Comparison of computational methods for Hi-C data analysis”. In: Nature Methods 14 (2017), pp. 679–685. doi: 10.1038/nmeth. 4325.

[35] Robert Tibshirani, Michael Saunders, Saharon Rosset, Ji Zhu, and Keith Knight. “Sparsity and smoothness via the fused lasso”. In: J. R. Statist. Soc. B 67 (2005), pp. 91–108.

[36] RJ Tibshirani and J Taylor. “The solution path of the generalized lasso”. In: Ann. Stat. 39 (3 2011), pp. 1335–1371.

[37] Trevor Hastie, Robert Tibshirani, and Martin Wainwright. Statistical learning with sparsity - the lasso and generalizations. CRC Press, 2015.

[38] Zheng Xu, Guosheng Zhang, Fulai Jin, Mengjie Chen, Terrence S. Furey, Patrick F. Sullivan, Zhaohui Qin, Ming Hu, and Yun Li. “A hidden Markov random field-based Bayesian method for the detection of long-range chromosomal interactions in Hi-C data”. In: Bioinformatics 32.5 (2016), pp. 650–656. doi: 10.1093/bioinformatics/btv650.

[39] Zheng Xu, Guosheng Zhang, Cong Wu, Yun Li, and Ming Hu. “FastHiC: a fast and accurate algorithm to detect long-range chromosomal interactions from Hi-C data”. In: Bioinformatics 32.17 (2016), pp. 2692–2695. doi: 10.1093/ bioinformatics/btw240.

[40] Holger Hoefling. A Path Algorithm for the Fused Lasso Signal Approximator. Vol. 19. 4. 2010, pp. 984–1006. doi: 10.1198/jcgs.2010.09208.

[41] Taylor B. Arnold and Ryan J. Tibshirani. genlasso: Path algorithm for generalized lasso problems. R package version 1.3. 2014.

[42] Bo Xin, Yoshinobu Kawahara, Yizhou Wang, and Wen Gao. “Efficient Generalized Fused Lasso and its Application to the Diagnosis of Alzheimer’s Disease.” In: AAAI. 2014, pp. 2163–2169.

[43] Álvaro Barbero and Suvrit Sra. “Fast Newton-type Methods for Total Variation Regularization.” In: ICML. Ed. by Lise Getoor and Tobias Scheffer. Omnipress, 2011, pp. 313–320.

[44] Álvaro Barbero and Suvrit Sra. “Modular proximal optimization for multidimensional total-variation regularization”. In: (2014). arXiv: 1411.0589.

[45] W Tansey and JG Scott. “A fast and flexible algorithm for the graph-fused lasso”. In: (2015). arXiv: 1505.06475.

[46] François Serra, Davide Baù, Guillaume Filion, and Marc A. Marti-Renom. “Structural features of the fly chromatin colors revealed by automatic three-dimensional modeling.” In: bioRxiv (2016). doi: 10.1101/036764.

[47] Stefan Lang and Andreas Brezger. “Bayesian P-Splines”. In: J Comp. Graph. Stat. 13.1 (2004), pp. 183–212.

[48] Stefan Lang and Andreas Brezger. “Generalized structured additive regression based on Bayesian P-splines”. In: Comput. Stat. Data Anal. 50 (2006), pp. 967–991.

[49] Andrew Gelman. “Prior distributions for variance parameters in hierarchical models (comment on article by Browne and Draper)”. In: Bayesian Anal. 1.3 (Sept. 2006), pp. 515–534. doi: 10.1214/06-BA117A.

[50] Carpenter B., Gelman A., Hoffman M., Lee D., Goodrich B., Betancourt M., Brubaker M. A., Gua J., Li P., and Riddell A. “Stan: A probabilistic programming language”. In: J. Stat. Software (in press).

[51] Stan Development Team. RStan: the R interface to Stan. 2016. url: http://mc-stan.org.

[52] JA Nelder and RWM Wedderburn. “Generalized Linear Models”. In: J. R. Statist. Soc. A 135 (1972).

